# Advanced 3D spheroid-based skin models and deep-learning based image analysis enable in-depth investigation of keratinocyte differentiation and barrier function

**DOI:** 10.64898/2026.01.21.700040

**Authors:** Tiziana Cesetti, Claudia Buerger, Nathalie Couturier, Elina Nürnberg, Roman Bruch, Victoria Lang, Mathias Hafner, Markus Reischl, Torsten Fauth, Rüdiger Rudolf

## Abstract

To date, a panel of different biological models have been used in skin research, ranging from *in vivo* testing to 2D and 3D cultures. Among these, organotypic skin models represent the current gold standard for preclinical dermatology and toxicology studies. However, they are variable in quality and require long maturation times and a lot of work and cells, the latter often of primary origin. Here, we propose dermal-epidermal spheroids as an alternative model that balances physiological relevance and throughput. Next to corresponding full thickness skin models, different fibroblast/keratinocyte coculture spheroids were generated. These used the commonly employed HaCaT cells as well as two recently immortalized keratinocyte cell lines, NHK-SV/TERT and NHK-E6/E7. To investigate their differentiation with detailed spatio-temporal resolution, a deep-learning segmentation-based pipeline, capable of revealing nuclear morphology and positioning as well as marker expression with single-cell precision, was developed and applied. Moreover, the formation of a functional barrier was assessed by live-imaging of Lucifer yellow diffusion. These experiments identified the NHK-E6/E7 cell line as the most and HaCaT cells as the least suitable alternative to primary keratinocytes in both spheroids and full thickness models. Furthermore, NHK-based coculture spheroids displayed functional maturation, including stratification, cornification, and barrier formation, closely recapitulating these features of corresponding full thickness models. Given the scalability and compatibility with automation, these micro-skin fibroblast/NHK-based 3D coculture spheroids might represent a promising new platform for pharmaceutical, cosmetic, and toxicological testing.

## INTRODUCTION

The skin, the largest organ of the human body, serves as a dynamic interface with the environment. Its complex structure fulfills essential functions including sensory perception, thermoregulation, and barrier formation, to protect the body from UV irradiation, dehydration, and chemical and microbial insults. To maintain these functions, the epidermis, skin’s outermost layer, is continuously renewed by proliferation of keratinocytes in the basal layer and their orderly differentiation and maturation into corneocytes in the apical layers. Each epidermal layer is characterized by the expression of specific markers: Cytokeratin 14 (KRT14) is primarily expressed in basal keratinocytes, marking the proliferative layer. Cytokeratin 10 (KRT10), a marker of early keratinocyte differentiation, is abundant from the lower spinous layer up to the granular layer, while involucrin (IVL) is expressed in the upper stratum spinosum and increases in the stratum granulosum ^[1]^. Filaggrin (FLG) is confined to the stratum granulosum, where it contributes to the formation of the cornified envelope and the skin barrier ^[1]^.

Beneath the epidermis lies the dermis, consisting of fibroblasts embedded in collagen and elastin fibers that provide strength and elasticity to the skin. The simplest human epidermal models, i.e., monolayer cultures of keratinocytes, are attractive due to their short cultivation time and low cost. Yet, they do not recapitulate the complexity of the multi-layered epidermis and many of its physiological functions. Animal testing, on the other hand, represents challenges in translating to human skin and raises ethical concerns. Thus, a variety of human 3D skin equivalent models have been developed over the past few decades and used for preclinical studies in dermatology, toxicology, and penetration testing ^[2,3]^. Simplified epidermal models can be generated by cultivating keratinocytes on an artificial substrate and inducing differentiation through exposure to the air-liquid interface (ALI) ^[4]^. In contrast, full-thickness skin models (FTSMs) are more advanced constructs including a dermal layer, consisting of matrix-embedded fibroblasts, overlaid by keratinocytes that mature into a stratified epidermis. When using primary keratinocytes isolated from human skin, reconstituted models closely resemble native skin, displaying a stratified and cornified epithelium and a functional barrier ^[5–7]^. However, primary keratinocytes show high donor-dependent variability, short life-span and limited availability ^[8,9]^. A common alternative is the spontaneously immortalized cell line HaCaT ^[10]^, which, however, exhibits only limited differentiation and is unable to form a functional epidermal barrier ^[11,12]^. To overcome these issues, several immortalized keratinocyte cell lines have been generated in the past ^[13–18]^ that better mimic characteristics of primary keratinocytes, also in the generation of 3D epidermal or full thickness skin equivalents ^[8,15,17,19]^. We have recently shown that the commercially available cell line NHK-SV/TERT (NHEK/SV-TERT3-5, Evercyte GmbH) ^[20,21]^ and our own cell line NHK-E6/E7 ^[22]^ are able to reconstitute a stratified epidermis and functional epidermal barrier in FTSM ^[23]^.

Despite this progress, *in vitro* skin models face several challenges, including their labor- and time-intensive manufacture, the need for a large amount of cells and the lack of robust automated procedures for either their generation and readout analysis ^[24]^. These limitations have restricted the use of *in vitro* skin models in high-throughput approaches, particularly impeding their implementation in early drug screening. Previously, self-assembling 3D cultures, termed spheroids, were introduced in cancer research as a compromise between physiological relevance and throughput ^[25]^. Spheroids can be easily and rapidly obtained in large quantities with homogeneous composition, size, and growth behavior ^[26]^. Typically, they are composed of some thousand cells with a few hundred microns in diameter in total. They can consist of a single cell type or different cell types, and are mostly superior in terms of physiological relevance compared to 2D adherent cell cultures ^[27,28]^. Usually, spheroid formation can be achieved by spontaneous cell aggregation in cell-repellent U-shaped 96-well plates ^[29,30]^, in agarose-coated plates ^[31,32]^, or hanging drops ^[33]^. Although their academic and industrial applications are rapidly expanding, especially in cancer research ^[34–39]^, the use of spheroids within the field of dermatology remains relatively limited ^[2,22,29,30,40,41]^. Thus, we aimed at establishing a new full-skin 3D culture system that is suitable for higher throughput, by generating coculture spheroids of fibroblasts with different keratinocyte lines. Their regular and reliable growth characteristics facilitated quantitative analysis. Therefore, we compared their stratification, morphology, and barrier function to classical *in vitro* skin-models in a quantitative manner. To this end, we developed a novel analytic tool set based on advanced image analysis and deep learning. Our data indicate that spheroids made of fibroblasts and either NHK-SV/TERT or NHK-E6/E7 cells easily form and that they exhibit a clear stratification, formation of tight junctions and a functional barrier, with NHK-E6/E7 spheroids performing best. Together, these factors might arguably render such spheroids a novel dermatological test model with intermediate complexity, physiological relevance, and throughput.

## RESULTS

### Generation of full thickness skin models and micro-skin spheroids

We aimed at developing a spheroid-based dermal-epidermal skin model with different keratinocyte lines and at evaluating their physiological significance by comparing them to corresponding ALI-based full thickness models (FTSM) in terms of differentiation and barrier formation. Spheroid cocultures were generated in ULA plates, while FTSM were produced by air-liquid interface culture (ALI) ^[42]^. In both systems, human CCD-1137Sk foreskin fibroblasts were first seeded, followed after 3 days (Figure 1A) by the addition of keratinocytes of three different cell lines, either NHK-SV/TERT, NHK-E6/E7, or HaCaT. While FTSM were air-lifted, spheroids were cultivated submerged in culture media (Figure 1A). HE staining showed that all FTSM formed an organized epidermis with signs of stratification and a dermal-like fibroblast layer of varying thickness. NHK-SV/TERT and NHK-E6/E7 in FTSM showed a higher degree of differentiation and stratification with an obvious outermost corneal layer (Figure 1B). Conversely, keratinocytes in HaCaT FTSM were not well organized and, particularly, showed no layers with granular or cornified appearance (Figure 1B). In NHK-SV/TERT and NHK-E6/E7 spheroids, a fibroblast core could be clearly distinguished at all time-points from the outer keratinocyte layers because of their different nuclear morphologies (Figure 1C). In contrast, the fibroblast nuclei in HaCaT spheroids appeared smaller at day 2 and were almost disappearing at later stages (Figure 1C). Fittingly, NHK-SV/TERT and NHK-E6/E7 spheroids maintained or slightly increased their size during the observation period, whereas HaCaT spheroids progressively decreased in size (Figure 1C). Altogether, this suggested the occurrence of cell death, primarily of fibroblasts and maybe also of keratinocytes, in the HaCaT, but not in the other co-culture spheroids. In support of this, the area of fibroblasts and of keratinocytes decreased over time in HaCaT spheroids, while it was constant or even increased in NHK-SV/TERT or NHK-E6/E7 spheroids (see below). All spheroids showed a denser peripheral layer of cells; however, internal stratification was less evident in HaCaT than in NHK-SV/TERT or NHK-E6/E7 spheroids, especially at D4 and D6 (Figure 1C). In particular, NHK-E6/E7 spheroids showed more distinct cell layers, with small well-defined cells close to the fibroblasts, and progressively flatter cells towards the periphery. By day 6, a densely packed outer layer with a few nuclei was suggestive of a stratum corneum (Figure 1C). In summary, both in FTSM and spheroids, NHK-SV/TERT and NHK-E6/E7 keratinocytes reconstituted a multilayered structure suggesting stratification and even cornification, while this was not evident with HaCaT cells.

**Figure 1:**
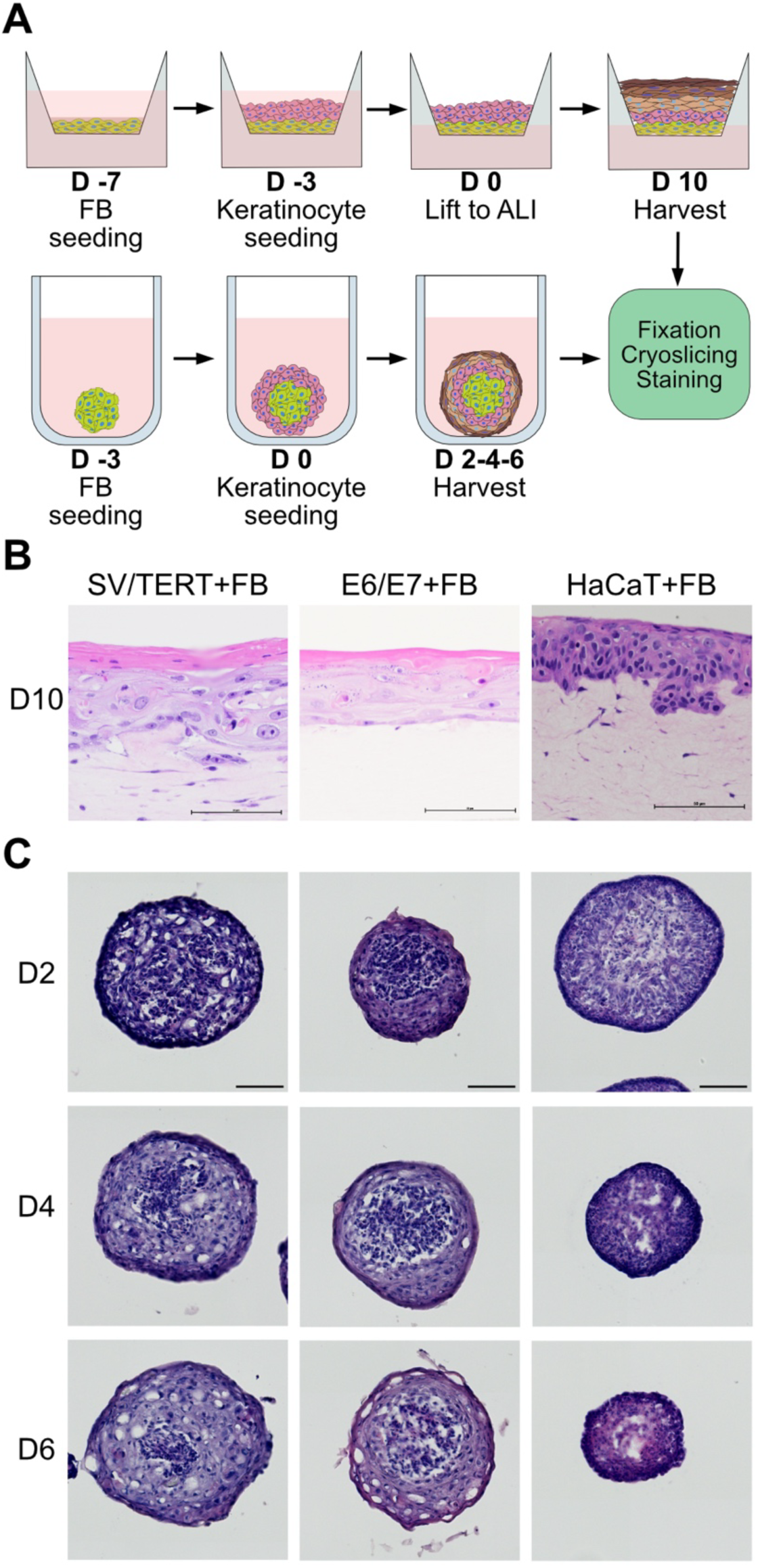
Morphological stratification is observed in both FTSM and spheroid-based dermal-epidermal cell models. **A)** Schematic illustration of the generation of FTSM (upper panels) and spheroids (lower panels). Initially, fibroblasts were seeded, either on collagen-coated transwell inserts (FTSM) or in ULA plates (spheroids). This was followed by addition of keratinocytes, either 4 days (FTSM) or 3 days (spheroids) later. FTSM were air lifted after 3 additional days and then grown at the air-liquid interface (ALI) for 10 more days before harvesting. Spheroids were cultivated submerged for 2, 4, or 6 days before harvesting. **B-C)** Representative H&E profiles of FTSM paraffine slices **(B)** and spheroid cryoslices **(C)**. Scale bars for FTSM and spheroids, 50 µm and 100 µm, respectively.

### Cellular differentiation and stratification in FTSM

In order to investigate differentiation and stratification of keratinocytes in FTSM, the expression of involucrin (IVL), cytokeratin 10 (KRT10), claudin 1 (CLD-1), and filaggrin (FLG) was analyzed by immunofluorescence upon cryosectioning (Figure 2A). While KRT14 was mainly detected in the basal layer in all models, KRT10 and IVL only showed correct localization in NHK-SV/TERT and NHK-E6/E7 FTSM. In contrast, both markers showed an overlapping expression across the epidermis in the HaCaT model. Additionally, FLG was detected in a broad peripheral band (Figure 2A). In NHK-SV/TERT and NHK-E6/E7 skin equivalents the tight junction marker claudin-1 (CLD-1) showed a nice cell-membrane staining in an intermediate region, as expected for tight junction formation in the stratum granulosum (Figure 2A). Conversely, in HaCaT FTSM, CLD-1 staining was more diffuse and not confined to the cell borders, indicating no or incomplete barrier formation (Figure 2A). In summary, these data suggested a superior epidermis-like differentiation and stratification in the FTSM produced with NHK-SV/TERT and NHK-E6/E7 than those made with HaCaT cells. As reduced differentiation often goes along with increased proliferation, it was fitting that HaCaT FTSM were overall much thicker (Figure 2B). In the epidermis, fully differentiated keratinocytes displayed flattened nuclei whose main axis is oriented in parallel to the border of the stratum corneum ^[43,44]^. To study keratinocyte nuclear morphology, we developed an automated analysis pipeline, based on deep-learning segmentation, in order to quantify nuclear shape and orientation. Quantification of nuclei features revealed a significantly reduced eccentricity of HaCaT nuclei compared to the more elongated nuclei of the other cell lines (Figure 2D). Furthermore, while the nuclei of NHK-SV/TERT or NHK-E6/E7 keratinocytes were aligned in parallel to the culture surface (lower values of relative orientation), such orientation was absent in HaCaT nuclei (Figure 2E). In summary, our data confirm previous observations ^[23]^, proving at a quantitative level the superior quality of skin models made from immortalized NHK cell lines compared to HaCaT in terms of stratification and differentiation.

**Figure 2:**
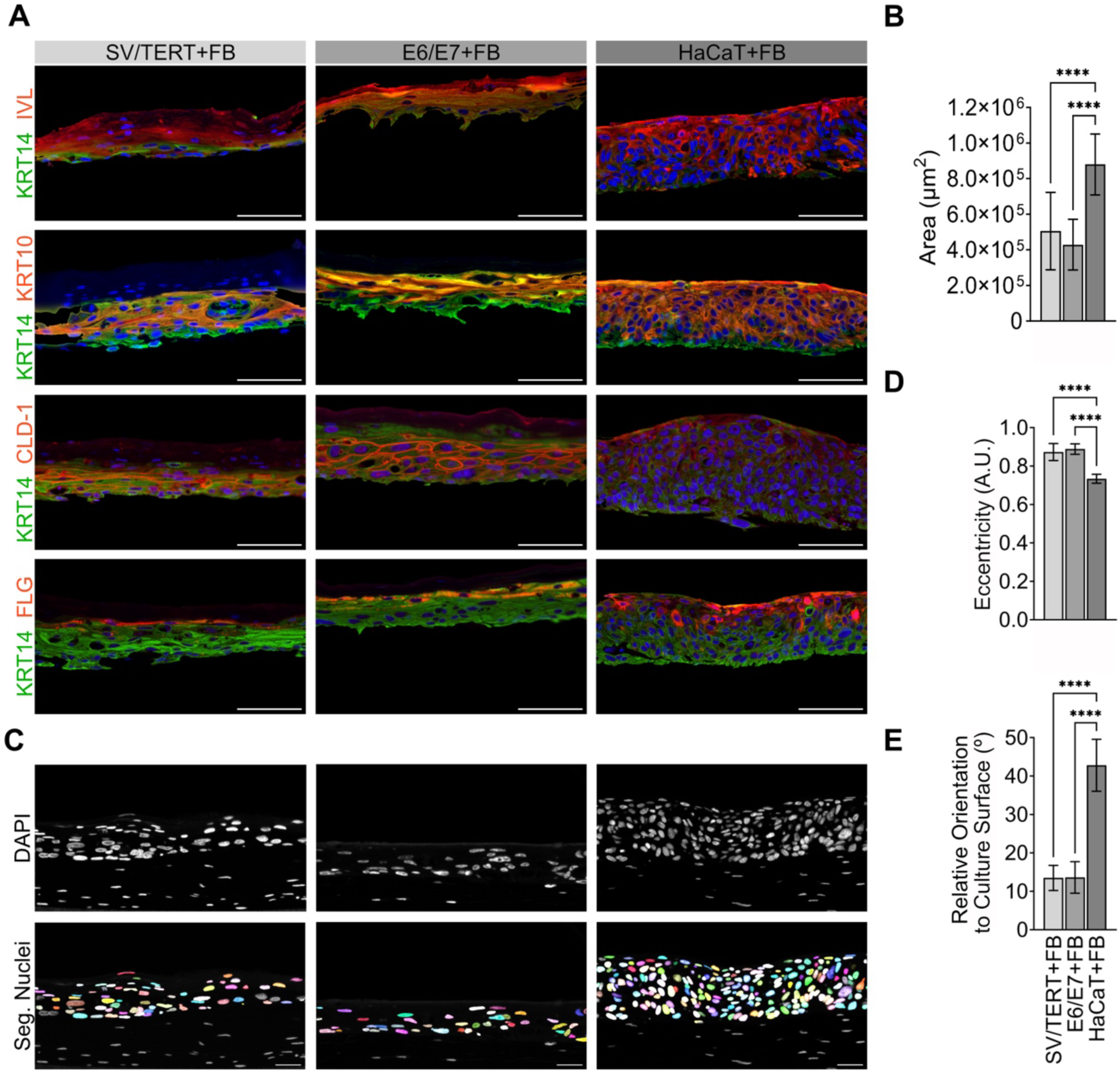
FTSM generated from NHK-E6/E7 or NHK-SV/TERT keratinocytes show stratification and differentiation, which is largely missing in HaCaT models. FTSM composed of fibroblasts and NHK-SV/TERT, NHK-E6/E7, or HaCaT keratinocytes were grown, harvested 10 days post ALI, fixed, sliced and stained for different protein markers, i.e., KRT14, IVL, KRT10, CLD-1, and FLG, as well as DAPI. This was followed by confocal microscopy. **A)** Representative images of FTSM upon immunostaining, DAPI shown in blue, other markers and their color codes as indicated. For a better visualization of the keratinocytes, the fibroblast regions were omitted. Scale bars, 50 µm. **B)** Plot of the area of the epidermal region. The keratinocyte region was defined based on the KRT14 signal for the lower border, and LY (not shown in this image) for the upper border. **C)** Representative images of nuclei (upper panels, DAPI) and overlay with the colored segmentation masks of keratinocyte nuclei (lower panels). Scale bars, 20 µm. **D)** Bar chart of keratinocyte nuclei eccentricity. Mean ± SD (N = 4 experiments). For a value close to 1, the nuclei are elliptic, lower values describe more roundish nuclei. **E)** Graph depicting the orientation of the nuclei long axis relative to the upper FTSM border. Mean ± SD (N = 16 experiments). A small angle indicates a parallel orientation of the nuclear long axis to the outer border. Two-way ANOVA in B, one-way ANOVA in D and E. **** p < 0.001.

### Barrier function in FTSM

Next, by a Lucifer Yellow (LY) penetration assay, we tested the functionality of a stratum corneum-like barrier in our FTSM. In the presence of a proper barrier, LY slightly penetrates only between corneocytes but not into the deeper layers. As depicted in Figure 3A, LY was retained in the upper regions of NHK-SV/TERT and NHK-E6/E7 FTSM, but not in those with HaCaT cells, in agreement with previous observation ^[8,12,19]^. Usually, LY images are shown as a proof of the presence of a functional barrier. However, differences may occur in the penetration depth. Therefore, we aimed at quantifying LY-staining characteristics by developing an image analysis routine. Plot profiles of LY intensity were calculated in the keratinocyte region (Figure 3A-B). This gave quantitative information on the depth and extent of dye penetration, which could be related to the structure and functionality of the different keratinocyte layers. Thus, a deeper penetration of LY was detected in NHK-SV/TERT than in NHK-E6/E7 FTSM. Indeed, plot profile analysis showed on average a deeper localization of the peak intensity (Figure 3C), a broader penetration window (Figure 3D), and a higher LY total intensity (Figure 3E) in NHK-SV/TERT than in NHK-E6/E7 FTSM. HaCaT FTSM did not show any LY signal, as the dye was not retained upon the wash-out, therefore they were not considered for the quantitative analysis (Figure 3B). In summary, these results suggest that the two NHK lines developed a functional barrier, but not HaCaT FTSM. Furthermore, significant differences in LY diffusion were detected between NHK-SV/TERT and NHK-E6/E7 FTSM, which might reflect distinct thicknesses and structures of the cornified and granular layers.

**Figure 3:**
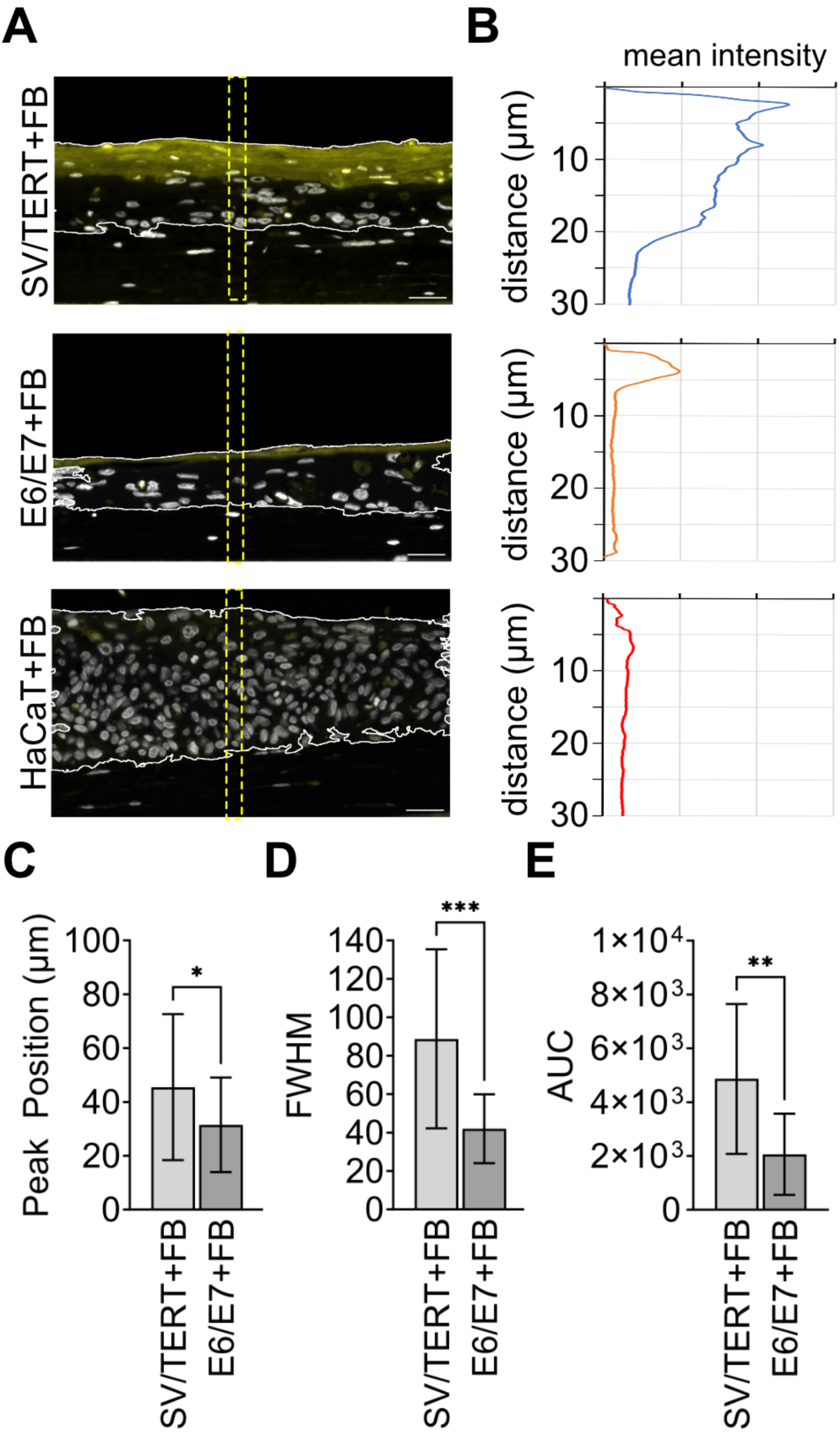
Barrier function is present in FTSM with NHK-E6/E7 and NHK-SV/TERT keratinocytes but not with HaCaT. FTSM composed of fibroblasts and NHK-SV/TERT, NHK-E6/E7, or HaCaT keratinocytes were grown, harvested 10 days post ALI, exposed for 1 h to LY, washed and then fixed, sliced and stained against nuclei (DAPI). **A)** Representative images of FT models after LY incubation, fixation, and slicing. Nuclei, gray; LY, yellow. Scale bars, 20 µm. **B)** Plot profiles depicting the mean intensity of the LY signal versus the depth (distance from the upper border), measured in the rectangular yellow dashed regions indicated in A. **C-E)** Quantitative analysis of LY staining results. Graphs show mean values ± SD for LY fluorescence peak position (distance from the upper border) **(C)**, full-width half maximum (FWHM) **(D)**, and the area under the curve (AUC) of the plot profile (all, N = 16) **(E)**. Significance was assessed with Mann-Whitney test for peak position and unpaired t-test for FWHM and AUC. The HaCaT FTSM model is not included in the graphs, since these showed no visible LY signal.

### NHK-spheroids reproduce important features of epidermal stratification

To gauge whether epidermal stratification occurs in fibroblast/keratinocyte spheroids, we performed comparable immunostaining experiments on spheroid cryoslices at different time points. Similar to what observed in Figure 1B, immunofluorescence images confirmed the presence of a fibroblast core surrounded by a keratinocyte region (Figure 4). A quantitative analysis of the spheroid total areas (see Figure 4 for technical description) confirmed that HaCaT spheroids were decreasing over time in size, while NHK-SV/TERT spheroids became larger and NHK-E6/E7 spheroids remained stable (Figure 4B). Looking at the individual compartments, the dermal core decreased over time in HaCaT spheroids, remained stable in NHK-SV/TERT and increased in NHK-E6/E7 (Figure 4C). The epidermal portion decreased in HaCaT, increased in NHK-SV/TERT, and was essentially unaltered in NHK-E6/E7 (Figure 4D). Overall, only HaCaT spheroids showed apparent loss of cells in both compartments.

**Figure 4:**
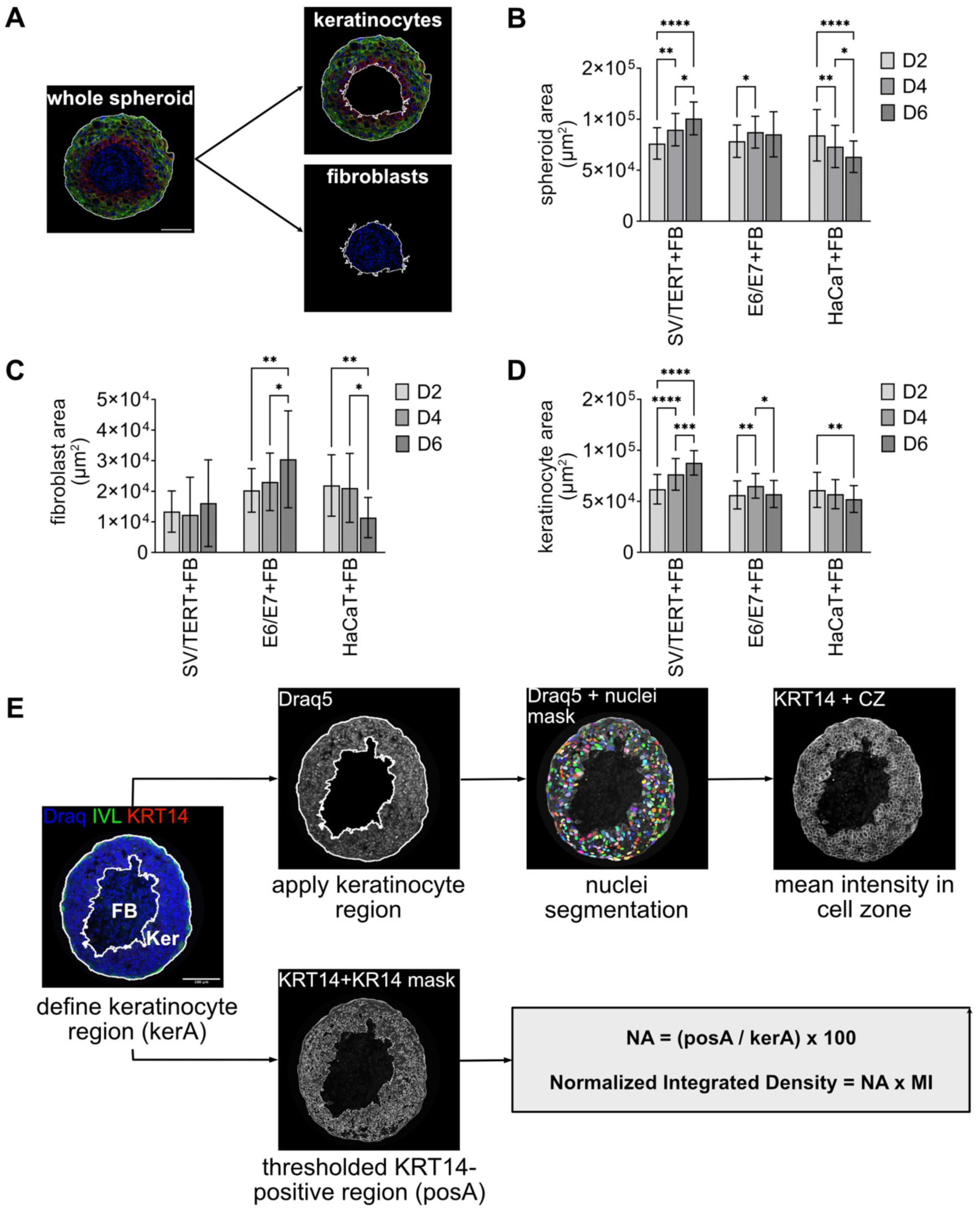
Molecular stratification is well developed in fibroblast-keratinocyte spheroids made with NHK but not with HaCaT keratinocytes. Spheroid coculture models composed of fibroblasts and NHK-SV/TERT, NHK-E6/E7, or HaCaT keratinocytes were grown and then harvested on days 2, 4, and 6 post coculture. Subsequently, spheroids were washed, fixed, sliced, and stained against nuclei (DAPI), KRT14, and IVL. **A)** Representative image of a spheroid cryoslice immunostained for KRT14 (red) and IVL (green). Nuclei, blue. The white borders indicate the regions selected as indicated. The areas of these regions were calculated with Fiji and the mean values ± SD were plotted of **B)** total spheroid area, **C)** fibroblast area, **D)** keratinocyte area. Statistics used two-way ANOVA (Turkeýs multiple comparison test) comparing for each cell line the different days after spheroid formation (D2, D4, D6) (N = 40). **E)** Schematic representation of the procedure to obtain segmented keratinocyte nuclei. Images were preprocessed to define and apply the keratinocyte region (to eliminate the fibroblast nuclei). Then, the nuclei were segmented with CellPose and masks were applied to the other channels to measure the mean intensity for each cell. In addition, the KRT14-positive keratinocyte region was used to measure the Normalized Integrated Density.

To get a deeper insight into the differentiation features and progression of the three spheroid models, a quantitative analysis was performed for the markers KRT14, KRT10 (Figure 4A, both red), IVL, FLG, and CLD-1 (Figure 5A, all green) over three observation time points.

**Figure 5:**
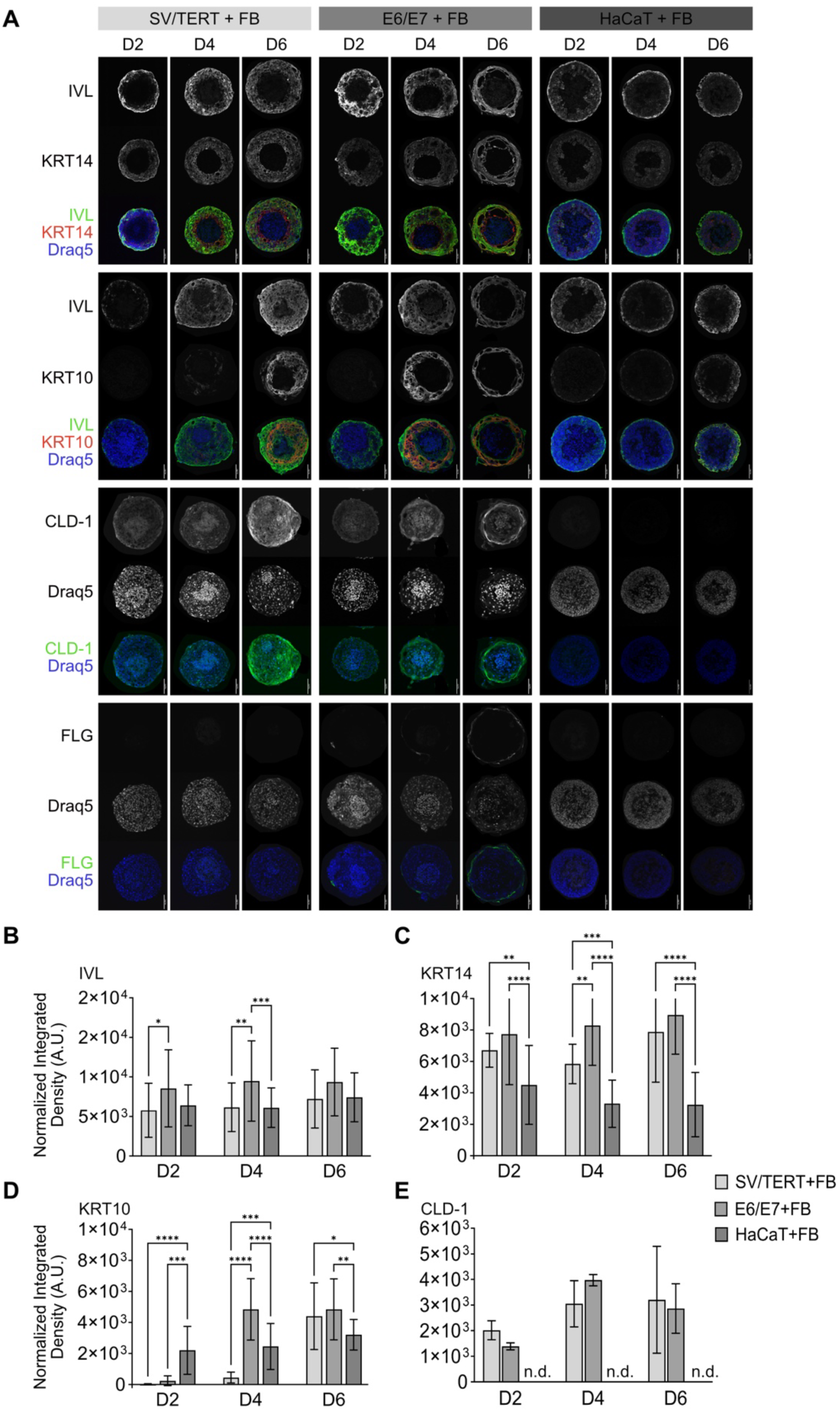
NHK-E6/E7 and NHK-SV/TERT spheroids show well-developed molecular stratification and mature over time, while HaCaT coculture spheroids exhibit weak stratification and deteriorate. Spheroid coculture models composed of fibroblasts and NHK-SV/TERT, NHK-E6/E7, or HaCaT keratinocytes were grown and then harvested on days 2, 4, and 6 of coculture. Subsequently, spheroids were washed, fixed, sliced, and stained against nuclei (Draq5) as well as for markers IVL, KRT14, KRT10, CLD-1, and/or FLG. **A)** Representative confocal images of cryoslices. Single-channel images of immunostained markers depict fluorescence intensities in gray, as indicated. Merge panels use color codes as indicated. Scale bars, 100 µm. **B-E)** Graphs of the normalized mean integrated densities ± SD at days 2, 4, and 6 for indicated markers. Statistics used Tukeýs multiple comparison test comparing, for each day, the 3 different cell lines (N between 20 and 30).

Confocal microscopy and data analysis revealed several notable differences between spheroids of different keratinocyte lines, as well as along the culturing time. While KRT14, KRT10, and IVL were detected in all types of spheroids, CLD-1 and FLG showed proper expression and distribution only in NHK-E6/E7 spheroids (Figure 5A). Their expression increased over time and became most evident at day 6, with FLG located in the outer spheroid rim and CLD-1 just beneath (Figure 5A), similar to the observation in FTSM (Figure 2A). In HaCaT spheroids KRT10 and IVL were present in the whole keratinocyte region, but enriched in the outermost zone. In contrast, in spheroids made from NHK-E6/E7 or NHK-SV/TERT cells, these markers showed a distinct distribution being essentially absent in the layer of KRT14-positive cells adjacent to the fibroblast core (Figure 5A). While IVL extended up to the spheroid border, KRT10 was mainly present in a central region and missing in the outer spheroid rim (Figure 5A). Quantitative analysis was performed by calculating the normalized mean integrated density, which considers both the mean intensity and the area covered by the signal (Figure 4E). This showed that, while IVL did not vary strongly over time and only little between the spheroid types (Figure 5B), KRT14 was much less present in HaCaT spheroids as compared to both other spheroid types (Figure 5C). Of all samples, NHK-E6/E7 spheroids displayed the highest expression of both IVL and KRT14 at all time points, suggesting that they have well-defined basal and differentiated keratinocyte layers. Interestingly, while KRT10 remained essentially unaltered at all time points in HaCaT spheroids, it markedly increased over time in the other spheroids, being earlier expressed in NHK-E6/E7 (day 4) compared to NHK-SV/TERT (day 6) (Figure 5D). Finally, CLD-1 was measurable only in the NHK keratinocyte spheroids, where it showed a trend towards increased expression from days 2 to 6 (Figure 5E). Overall, these data suggest a wide gap between the NHK lines and HaCaT spheroids. Similar to what was observed in FTSM, NHK-E6/E7 spheroids showed the best recapitulation of epidermal differentiation and stratification, particularly as suggested by the expression pattern of KRT10, CLD-1, and FLG.

### Single-cell analysis of nuclear features and marker expression in spheroids

The previous analysis has shown that NHK-E6/E7 and NHK-SV/TERT keratinocytes in spheroids have a higher and more stable expression of differentiation markers compared to HaCaT. However, regional differences and features of cell morphology were not taken into account. To better describe the stratification and the ongoing differentiation process, we therefore aimed at gaining information at a single-cell level by linking marker expression with cell position and nuclei features. For this purpose, we developed a single-cell analysis workflow. First, keratinocyte nuclei were automatically segmented using a deep-learning algorithm (Cellpose) (Figure 4A), then a program code was developed to retrieve nuclei orientation versus the outer spheroid border and nuclei eccentricity. Finally, these features were related to the cells’ positions within the spheroid, obtained by measuring nuclei distance to the outer rim and to the keratinocyte/fibroblast interface. The keratinocytes were then segregated into three different regions, based on their relative position: a surface region (R1), a core region (R3), and an intermediate region (R2) (see Material and Methods). Regarding eccentricity, single-cell analysis showed that on day 2 in all spheroid types the nuclei located in R1 had a significantly higher mean value compared to the other regions, indicating a more elongated shape (Figure 6B). This proves an adaptation of keratinocyte morphology, with the outer cells displaying more elongated nuclei, a feature of terminal differentiation that we similarly observed in NHK-FTSM (Figure 3D). This difference in eccentricity persisted in NHK-E6/E7 and NHK-SV/TERT spheroids over time, but was lost in HaCaT spheroids after day 2, suggesting a loss of differentiated cells at the later time points. These findings were corroborated by the analysis of nuclear orientation. In NHK spheroids, for all time points analyzed, the nuclei in R1 were more aligned to the outer border than those in R2 and R3 (Figure 6C). Conversely, in HaCaT spheroids, R1 nuclei showed a significant alignment difference only on day 2 (Figure 6C).

**Figure 6:**
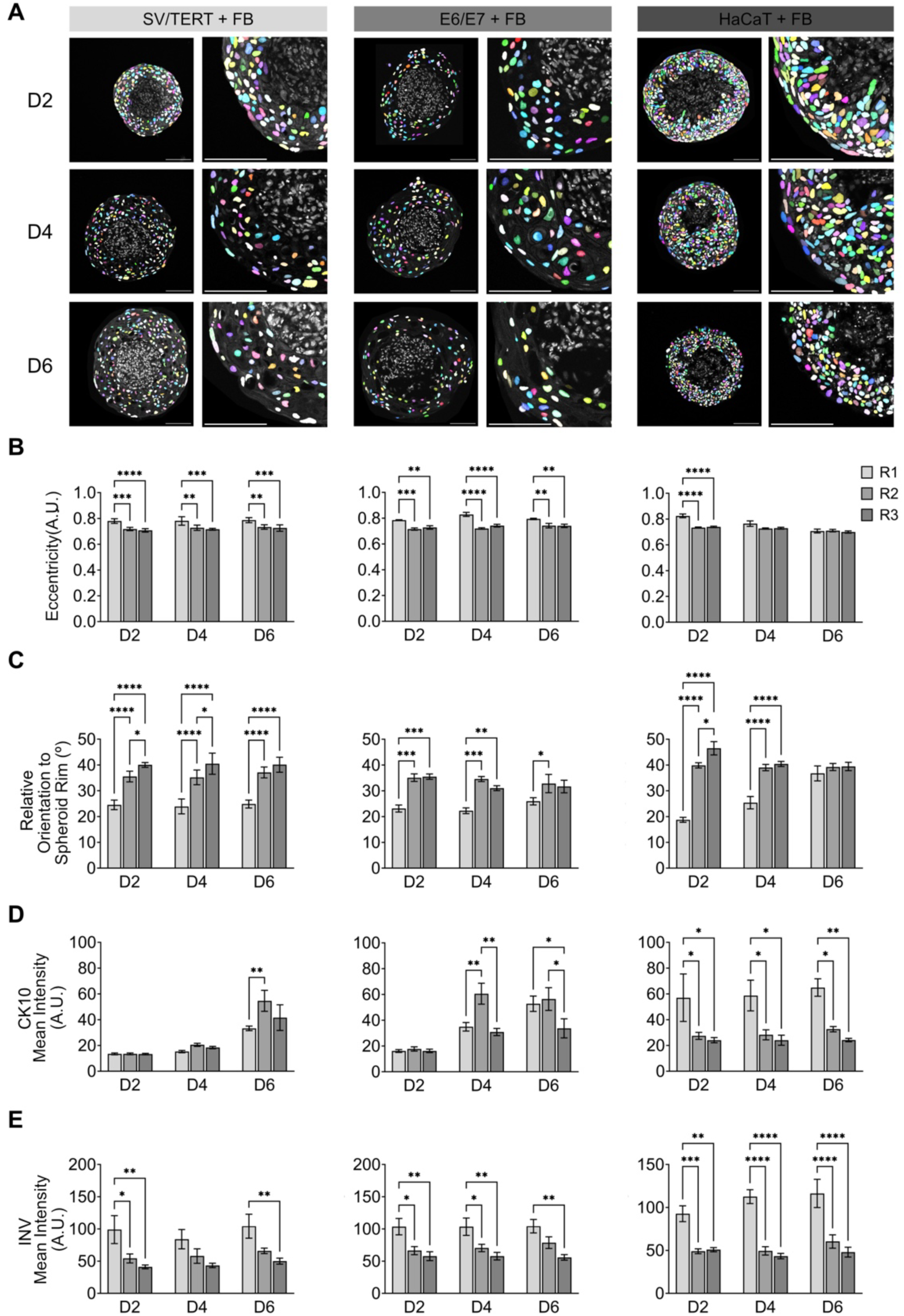
Single-cell analysis of nuclei features and marker localization confirm superior stratification of NHK-E6/E7 and NHK-SV/TERT over HaCaT coculture spheroids. Spheroid coculture models composed of fibroblasts and NHK-SV/TERT, NHK-E6/E7, or HaCaT keratinocytes were grown and then harvested on days 2, 4, and 6 of coculture. Subsequently, spheroids were washed, fixed, sliced, and stained against nuclei (Draq5) as well as for markers, IVL and KRT10. Keratinocyte nuclei or marker intensities were automatically segmented and quantified. **A)** Segmentation masks of keratinocyte nuclei (colored) were superimposed onto Draq5 raw images (gray). For each cell line, confocal images of spheroids taken at days 2 (D2), 4 (D4), or 6 (D6) were taken. Left panels show representative overview images, right panels their higher magnification. Scale bars, 100 µm. **B-E)** For each spheroid, the segmented nuclei and cell zones were grouped based on their position. R1: 0 - 20 µm from the outer spheroid border. R3: 0 - 20 µm from the fibroblast core. R2: between R1 and R3. Bar plots represent the mean of 4 technical replicates ± SD. Each technical replicate comprehends 5-10 spheroids, each spheroid slice has 100-200 cells. Statistics used two-way ANOVA comparing R1, R2, and R3. **B)** Graphs of nuclear eccentricity: higher values represent more elliptic nuclei. **C)** Graphs of nuclei relative orientation: lower values represent nuclei parallel to the outer border. **D, E)** Graphs of the mean intensity in the cell zone for KRT10 **(D)** or IVL **(E)**.

The single-cell analysis with regional clustering was also used to probe stratification of differentiation markers. For HaCaT spheroids, both IVL and KRT10 expression were strongest in the R1 cells and this was constant over time (Figure 6D and E). In contrast, KRT10 was almost negligible at day 2 in the other two spheroid types. Its expression was first observed on days 4 in NHK-E6/E7 and on day 6 in NHK-SV/TERT spheroids (Figure 6D and E). Notably, KRT10 was concentrated in the middle region (R2), similarly to FTSM. Finally, IVL was mostly enriched in R1 of all spheroids at all time points (Figure 6E), but at difference with HaCaT, it showed a more gradual decrease toward the spheroid core in NHK-E6/E7 and NHK-SV/TERT spheroids (Figure 6E). In summary, these data further support the finding that immortalized-keratinocyte spheroids are superior to HaCaT spheroids in terms of epidermis-like organization and stratification.

### NHK spheroids exhibit a functional barrier

To investigate a final differentiation to corneocytes and the formation of a functional barrier in spheroids, similar to what was observed in FTSM, we built upon our previous experience with live spheroid confocal imaging under perfusion ^[45]^ to set up a new protocol that allows time-course measurements of spheroids while applying LY (Figure 7). In this configuration, while LY penetration should be precluded by an existing barrier, it may diffuse inside the spheroid and be retained there if the barrier is absent. In order to define the spheroid region, we took advantage of the cell-permeant dye, CellMask (CM, Figure 7A). The live labelled spheroids were imaged at three different z-depths (0, 35, and 70 µm), to evaluate the capability of LY to diffuse into deeper spheroid regions. Spheroids were first perfused with a control solution (-LY), and then with LY (+LY), followed by wash out (Figure 7A, B). LY perfusion led to a sudden increase of fluorescence in the solution surrounding the spheroid and, with a small delay, also within the spheroid. The fluorescence measured in the spheroid was normalized to the fluorescence outside and plotted over time (Figure 7B). This showed that in NHK-SV/TERT and NHK-E6/E7 spheroids, the change of LY fluorescence was larger at the outer part (0 µm) than in the deeper levels, while in HaCaT spheroids all levels showed a similar behavior (Figure B). This suggests that LY diffused deeper into HaCaT compared to NHK spheroids. At 35 and 70 µm depth, the amount of LY fluorescence was indeed significantly larger in HaCaT compared to the other two cell lines (Figure 7C). To corroborate a functional barrier formation, SDS pretreatments are usually employed in FTSM ^[5,19,46]^. Fittingly, SDS pretreatment abolished the differences regarding LY penetration between the three spheroid types at all depths, strongly arguing for a compromised barrier integrity in NHK spheroids under these conditions (compare Figures 7B and D).

**Figure 7:**
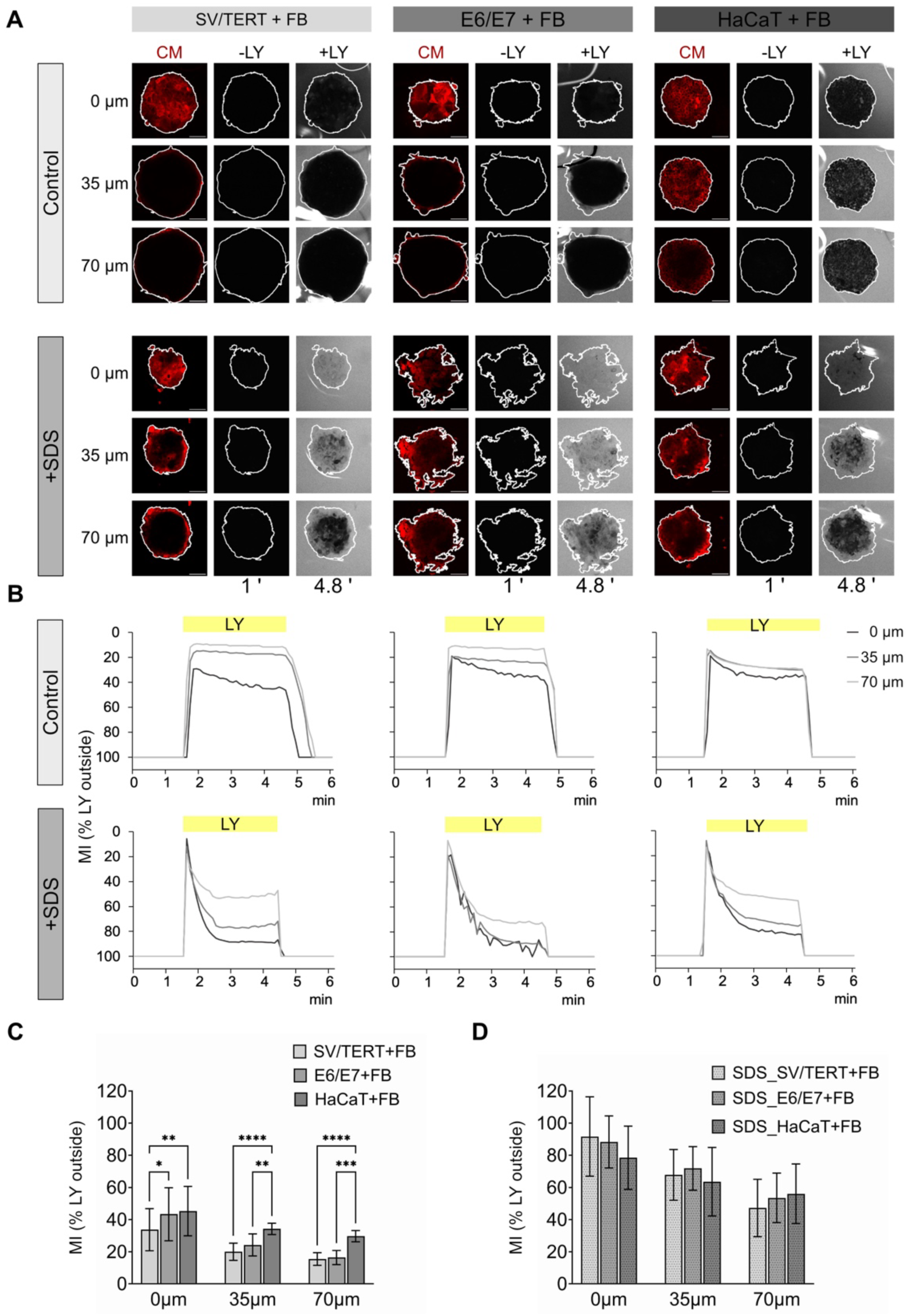
Lucifer Yellow diffuses deeper into coculture spheroids made with HaCaT compared to those with immortalized keratinocytes. **A)** Representative images in control or upon SDS pretreatment, at different depths from the spheroids’ outer borders, as indicated (0-35-75 µm). Left panels for each cell type: CellMask (CM) fluorescence, used to define the spheroid region (white outline). Central and right panels for each cell type: LY fluorescence in the absence (-LY) and in the presence (+LY) of the dye in the perfusion solution. Images were taken at the times indicated below the bottom panels. White outlines (CM positive) define the spheroid region. Scale bars, 100 µm. **B)** Time plots of LY fluorescence signal in the spheroid, at different depths, as indicated. The mean intensity was calculated in the spheroid region and normalized to the region outside the spheroid (MI (% LY outside)). In the absence of LY, the spheroid and the background are similarly dark (100 %). Upon LY perfusion, the outer region becomes brighter than the spheroid, thus the spheroid fluorescence is only a fraction of the background. **C, D)** Bar plots represent the mean of MI (% LY outside) ± SD, measured at the end of LY perfusion (4,8 min). Statistics used two-way ANOVA, comparing spheroids of different cell types for distinct depths and in the absence (**C**) or presence of SDS (**D**) (N = 10 - 22).

As LY diffused not only in depth, but also radially into the spheroids, we wanted to quantify the amount of LY fluorescence at the outer border compared to the inner spheroid region. For this purpose, the data were reanalyzed by defining an outer region for each spheroid, i.e., reflecting a potentially cornified zone, and a core region, i.e., including the rest of the keratinocytes and the fibroblasts (see Material and Methods). Upon LY perfusion, NHK-SV/TERT and NHK-E6/E7 spheroid core regions remained essentially devoid of LY signal, while LY diffused into the inner part of HaCaT spheroids (Figure 8A). Quantitative analysis confirmed that LY fluorescence in the spheroid core (MI_in_) was consistently higher in HaCaT as compared to the two NHK spheroid types (Figure 8D), while fluorescence in the outer region (MI_out_) was comparable between all spheroid types (Figure 8E). In summary, this led to a significantly higher MI_out_/MI_in_ ratio in NHK-E6/E7 or NHK-SV/TERT, as compared to HaCaT spheroids (Figure 8F). Notably, SDS pretreatments led to a massive diffusion of LY into the cores of all spheroid types and thus ablated any difference previously observed (Figure 8 G-L). In summary, LY live-imaging experiments showed that this hydrophilic dye was much more excluded from the cores of NHK-SV/TERT and NHK-E6/E7 spheroids, arguing for the existence of a functional barrier in these spheroids, similar to what observed in FTSM. This is also in agreement with previous observations in fibroblast/keratinocyte spheroids obtained either with HaCaT or primary keratinocytes, where, upon long LY incubation, only HaCaT spheroids were massively loaded ^[47]^. Our LY diffusion experiments complemented the marker expression analysis to prove that only in NHK spheroids the relevant structural, cellular, and biochemical components were present, which contributed to the generation of the functional barrier. Furthermore, our new experimental setting for LY diffusion was not only suitable to test the presence of a barrier, but it also provided additional spatial information and temporal kinetics for a more detailed study of the barrier properties and small differences therein.

**Figure 8:**
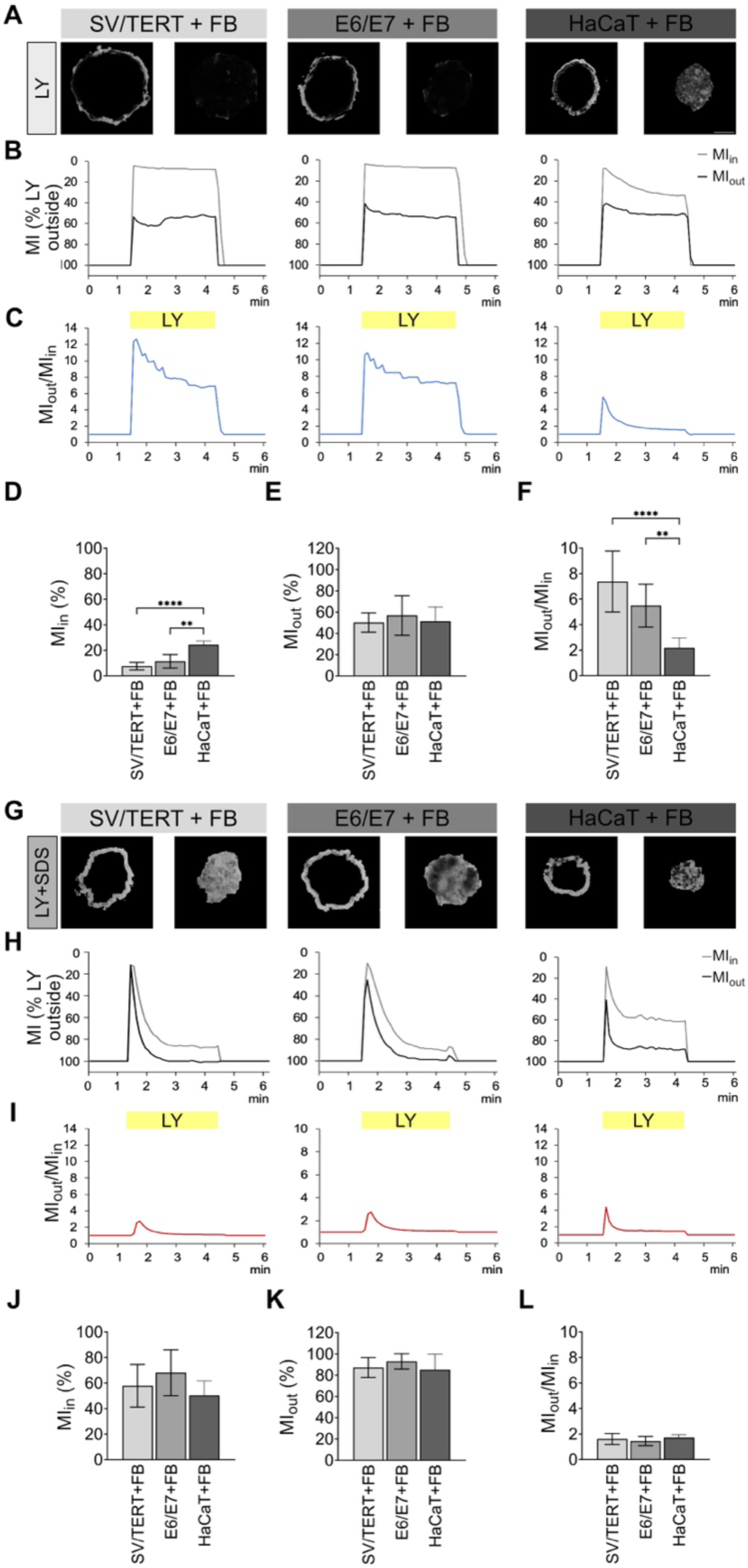
Lucifer Yellow diffuses into the core of HaCaT spheroids, but is hindered in the other spheroids. Spheroid coculture models composed of fibroblasts and NHK-SV/TERT, NHK-E6/E7, or HaCaT keratinocytes were grown for 4-6 days and then live-imaged with confocal time-lapse microscopy. During imaging, spheroids were perfused first in control and then with LY at the time indicated in the plots (yellow bars). Where indicated, they were preincubated with SDS. **A-F)** Perfusion experiment in the absence of SDS. Coculture type as indicated on top. **A)** Representative confocal images of coculture spheroids at a depth of 35 µm at 3 min after LY perfusion. Left and right panels of each image pair show only the LY fluorescence intensity in the spheroid border and center, respectively. LY fluorescence outside the spheroids was clipped for better visibility. For full images, see Figure 7. Scale bar, 100 µm. **B-C)** Graphs show time plots of representative experiments. LY mean intensities in the spheroid regions are depicted as percent of LY fluorescence outside the spheroid (MI (% LY outside)). Time plot of LY mean intensity at the spheroid border (MI_out_, dark gray) and in the center (MI_in_, light gray) **(B)** and of their ratio (MI_out_/MI_in_) **(C)**. **D-F)** Bar plots represent the means of MI (%) ± SD, as measured in the inner **(D)** and outer regions **(E)** and the mean of their ratio, MI_out_/MI_in_ **(F)**. **G-L)** LY-perfusion experiment upon SDS pretreatment. Coculture type as indicated on top. **G)** Representative confocal images of coculture spheroids at a depth of 35 µm at 3 min after LY perfusion. **H-I)** Graphs of LY mean intensities of spheroid border (MI_out_, dark gray) and center (MI_in_, light gray) as percent of outside LY fluorescence **(H)** and MI_out_/MI_in_ ratio **(I)**. Yellow bar, denoted LY, indicates perfusion period of LY. **J-L)** Bar plots, mean of MI% ± SD, as measured in the inner **(J)** and outer regions **(K)** and the mean MI_out_/MI_in_ **(L)**. Statistics used two-way ANOVA (B, C, F, G) or Kruskal-Wallis nonparametric test with Multiple comparison (D, H) (N = 13 - 17).

## DISCUSSION

This work showed that coculture spheroids with fibroblasts and NHK-SV/TERT or NHK-E6/E7 keratinocytes were capable of fully differentiating into a mature epidermal structure, similar to FTSM obtained with the same cells. Thus, we propose these spheroids as a promising alternative 3D skin model. Furthermore, we developed and applied new quantitative analysis tools which allowed a detailed comparison of keratinocyte maturation, stratification, and barrier function across both FTSM and spheroid formats.

For the immunostained cryoslices we developed an analysis pipeline that allows the quantification of marker expression and nuclear features at single-cell precision, and that combines this information with the position of the cells within the spheroids. Through this detailed quantitative analysis, we proved that NHK-SV/TERT and NHK-E6/E7 coculture spheroids displayed progressive maturation and stratification, in line with what we observed in comparable FTSM. On the contrary, HaCaT spheroids shrank over that time, underwent disorganization and did not fully maturate. A further sign of full epidermal maturation of NHK-SV/TERT and NHK-E6/E7 coculture spheroids was their presence of a functional barrier, as tested by the LY diffusion assay. Interestingly, NHK-based spheroids blocked LY penetration, while with HaCaT the dye diffused into the spheroid core. These data are in line with FTSM results and also with previous epidermal ALI-based skin models generated with the same cell lines ^[23]^. In summary, our data support the generation of a stratified epidermis and of a functional barrier in FTSM and spheroids composed of fibroblasts plus either NHK-SV/TERT or NHK-E6/E7, but not of HaCaT cells.

Thus, due to their matched morphological and functional features with FTSM, NHK coculture spheroids introduced here represent a miniaturized 3D skin model with distinct advantages over classical human skin equivalents. Their applications in the field of dermatological research may increase when considering the following benefits. First, the production can be scaled, considering the much lower number of cells required for spheroid generation compared to organotypic skin models. Secondly, they can be handled in multi-well plates, therefore enabling automated processing and reducing culture volumes. Third, given their rapid formation and differentiation, dermal-epidermal spheroids reduce the time required to generate corresponding skin equivalent cultures by at least one half, significantly saving time. Fourth, since spheroids are processed throughout in submerged culture, drug treatments should be easier to perform and require smaller amounts compared to the ALI cultures.

Next, regarding the use of HaCaT cells to generate 3D skin models, this study provides a detailed temporal and spatial analysis of differentiation, stratification, and functional barrier, which proves the similarity between spheroids and ALI-based skin model with HaCaT cells and extends what was already known from literature. Indeed, it was previously reported that organotypic skin cultures with HaCaT keratinocytes initially form a multilayered epithelium, which increasingly disorganizes and undergoes senescence ^[12,48]^. In particular, while KRT10 expression was restricted to the uppermost layer after one week in culture in skin equivalents, it was ubiquitously expressed at later time points ^[12]^. This was at difference with primary keratinocytes, showing KRT10 expression restricted to the suprabasal layers in FTSM ^[49]^. Similarly, our results in HaCaT-based FTSM show ubiquitous expression of KRT10 and the presence of lobulated and irregular nuclei also in the outermost layer, while in native skin the keratinocytes of the stratum corneum are flattened and anucleated ^[12]^. HaCaT easily form spheroids, either alone or with a fibroblast core ^[2,26,30,39,50]^. Fibroblast/HaCaT spheroids reproduced some epidermal features, including KRT10 and KRT14 expression, nevertheless KRT10 localization was incorrect ^[50]^. Additionally, complex stratification was not reported and barrier function has not been tested, so far ^[26,29]^. Our marker expression analysis in HaCaT coculture spheroids suggested an advanced differentiation state compared HaCaT monoculture spheroids, but still they were all lacking a correct stratification and the formation of a cornified layer. While at day two HaCaT nuclei shape and orientation suggested a kind of organization of the cell layers, HaCaT spheroids underwent increasing disorganization and cell death, which make this model not suitable for long cultivation experiments. Additionally, single-cell analysis clearly proved incorrect localization of KRT10, while the lack of FLG and CLD-1, and of a functional barrier proved the absence of a stratum corneum. Thus, our data clearly define the limitations of HaCaT based dermal/epidermal spheroids.

Conversely, some immortalized NHK cells were already shown to have a higher physiological relevance compared to HaCaT. Along these lines, another immortalized NHK line was previously reported to generate FTSM displaying a higher similarity to primary keratinocytes, with correct localization of KRT10 in the stratum granulosum and spinosum ^[51]^. In our case, FTSM generated with NHK-SV/TERT or NHK-E6/E7 displayed all four epidermal layers, according to marker expression. In particular, FTSM with NHK-E6/E7 displayed the best stratification and showed a marker distribution comparable to FTSM obtained with primary keratinocytes ^[23]^. Interestingly, the nuclear features, elongation and parallel orientation, spoke also in favor for layering and terminal differentiation in the two NHK-based FTSM.

Similar features were observed also in our NHK coculture spheroids. While IVL expression was more pronounced in the outer region and constant over time, the expression of the other differentiation markers increased at later time points, suggesting a progressive differentiation of the keratinocytes along the six days of cultivation. In particular, the single-cell analysis revealed that KRT10 expression was largely restricted to a central region, that could correspond to the suprabasal layer. Such localization was observed in both NHK-based spheroid types, but expression was delayed in NHK-SV/TERT. This confirmed a previous speculation that in NHK-SV/TERT the switch from proliferation to differentiation may take longer than in NHK-E6/E7 ^[23]^. A more stratified and differentiated phenotype in NHK-E6/E7 spheroids was also supported by the expression of CLD-1 which was better localized in a thin outer cell layer. Similarly, FLG was exclusively present in NHK-E6/E7-spheroids, where it was properly restricted to the outermost region, i.e., corresponding to the granular layer. This expression pattern was paralleled by a progressive flattening of the keratinocyte region and by elongation and alignment of the nuclei with the outer spheroid rim. Moreover, the presence of a functional barrier in NHK-spheroids was reinforced by the exclusion of LY in live-imaging experiments. This corroborates the occurrence of a terminal differentiation and suggests the formation of a competent stratum corneum, especially in NHK-E6/E7 spheroids. To our knowledge, this is the first evidence that a functional skin barrier can be achieved in a micro-skin spheroid model generated with keratinocyte cell lines, which highlight the similarity with primary keratinocyte spheroids ^[47]^.

Only a few groups have pursued the generation of spheroids from keratinocytes other than HaCaT. Notably, the presence of a fibroblast core was key to achieve multilayered spheroids ^[47,52]^. Primary keratinocytes, when cultivated in BME droplets, developed an inside-out epidermis model, with the corneocytes in the core ^[53]^. Spheroids could be generated with primary human neonatal keratinocytes, but with low efficiency and neither differentiation nor stratification were proved ^[32,54]^. In coculture spheroids, generated with iPSC-derived fibroblasts, keratinocytes, and endothelial progenitors, the keratinocytes organized around the fibroblast core, expressed KRT14 and KRT10, but stratification and cornification were not present ^[55]^. More successfully, micro-skin organoids, generated with a fibroblast/collagen core and keratinocytes derived from epiblast cells, fully differentiated and developed a functional barrier ^[47]^. In our case, NHK cells could not form spheroids without fibroblasts, but only loose cell aggregates. Instead, stable spheroids were formed in the presence of fibroblast-conditioned medium, but in the absence of a fibroblasts core, the keratinocytes did not show stratification (data not shown). Thus, in essence, our dermal-epidermal spheroid cultures with NHK-lines showed a higher level of cell differentiation and layering, adding physiological significance to spheroids as skin models. Particularly remarkable was the coexistence of basal and differentiating keratinocytes in the same spheroid but in different regions, with KRT14-positive cells mainly present around the fibroblast core, i.e., corresponding to the basal layer, and the other markers localized according to a physiological skin differentiation program.

There are several potential use-cases and further practical implications for the fibroblast/NHK spheroids and for the assays presented in this study. First, considering the applications of the confocal live-imaging LY assays, our experimental setting and kinetic analysis have allowed a detailed quantification of LY penetration and its time course. Our live-imaging study of diffusion in spheroids might open new ways to study permeation and absorption of drugs, which can be relevant to develop formulations for transdermal drug delivery ^[3,59]^. Several concentrations could be tested thorough the perfusion in a dynamic way. It has also potential for high-throughput applications, when using an automated confocal system. However, ULA plates are not suitable for confocal microscopy due to the thick plastic and concave shape, thus they should be substituted with imaging-compatible polymer arrays ^[63,64]^. The perfusion systems should be also automated, to allow defined and reproducible treatments. In a previous work, HaCaT-based dermal-epidermal spheroids were generated with hydrogels in a microfluid assay and used for screening the efficacy of vitamins and peptides and the toxicity of chemical agents. They were also tested for diffusion of CellTracker-Red-CMTPX, which labelled HaCaT cells, but did not reach the fibroblast core ^[41]^. This paper posed the basis for high-throughput screening using skin spheroids and live-imaging. Thus, a combination of our NHK-based spheroids with an automated confocal live-imaging and microfluidic system might allow the generation of a high-throughput platform to study skin barrier integrity and diffusion of fluorescently labelled molecules. NHK-based spheroids could complement pharmacologic, toxicologic and cosmetic tests that so far take place in human skin equivalents or in animals *in vivo*, opening the possibility of larger and faster screenings ^[57,58]^.

Second, considering the improved physiological significance of our NHK-based spheroids, they could be useful to model diseases. Previously, it was shown that incorporation of melanoma cells into fibroblast/HaCaT spheroids significantly reduced KRT10 enrichment at the spheroid rim. A cytostatic treatment efficiently killed melanoma cells and recovered the peripheral localization of KRT10 ^[29,30]^. It would be interesting to model melanoma in triculture spheroids with the immortalized NHK cells, in order to investigate how cancer cells would affect their differentiation and prove drug efficiency in recovering stratification and barrier function ^[56]^.

Of note, the conclusions in this study were not merely based on qualitative analysis, but achieved by detailed quantitative information. The presented advanced segmentation-based image analysis of immunostained cryoslices yielded quantitative single-cell information, i.e., describing for each cell its position, nuclear morphology and orientation, and marker expression. Notably, this novel analytical approach, when applied to both spheroids and FTSM, can allow a more precise spatial information. In particular, nuclei features could be used as a reliable read out of differentiation and stratification in the case of the skin. This approach could be used also to study differentiation and cell patterning in other tissues and organoids ^[60–62]^.

At present, though, there is a major bottleneck in terms of throughput for our experimental design. Indeed, immunofluorescence microscopy of both skin equivalents and spheroids relies on the tedious and time-consuming process of cryosectioning. Thus, to unleash the full performance of spheroid-based experimental studies, it would be important to circumvent the need for cryosectioning and instead enable whole-mount immunostaining and 3D analysis. This can be better multiplexed and automated. Previously, general protocols of whole mount immunostaining and optical tissue clearing of 3D cell cultures were devised to meet exactly this need ^[65,66]^. Yet, while the available methods were found to be applicable to HaCaT spheroids ^[65,67,68]^, they were ineffective on NHK-based spheroids (data not shown). This was most likely due to the enhanced barrier function of these spheroids, which did not permit the penetration of either antibodies, staining reagents, and/or light. New clearing protocols for skin spheroid models should be developed based on the results already obtained on skin biopsies ^[69–71]^. Future work should address this challenge in more detail to further improve the benefit and speed at which fibroblast/NHK-based spheroids can address open topics in experimental dermatology.

In summary, we developed spheroids consisting of fibroblasts and immortalized keratinocytes that matured in robust, stratified, barrier-competent dermal/epidermal structures. Together with our deep-learning-based image analysis tools, these models offer an intermediate-complexity, physiologically relevant, and moderately high-throughput platform for dermatological testing.

## MATERIALS AND METHODS

### Cell cultivation

<u>CCD-SK1137 Fibroblasts:</u> CCD-1137Sk human fibroblasts were from American Type Culture Collection (ATCC, Manassas, VA, USA); they were cultivated in IMDM supplemented with 10% FBS at 13.000 cells/cm^2^ density. <u>HaCaT keratinocytes:</u> HaCaT were from CLS Cell Lines Service GmbH (Eppelheim, Germany). They were cultivated in DMEM medium with Na-Pyruvate (1 mmol) and Glutamine (4 mmol) (SIGMA D5796), supplemented with 10% FBS. Cells were grown in plastic flasks (Greiner) at a density of 5000 cells/cm^2^. <u>NHK/SV-TERT3-5</u>: were purchased from Evercyte GmbH (Austria). They were obtained from human epidermal keratinocytes by transfection with a plasmid encoding SV40 early region followed by transduction with a retroviral expression vector (pLXSN) containing the TERT gene. <u>NHK-E6/E7</u>: were provided by BRAIN Biotech AG (Germany). Juvenile keratinocytes were immortalized by lentiviral transfection with HPV16-NHK-E6/E7. The two latter cell lines were cultivated in 2D-KGM medium (Promocell) at a density of 6000-8000 cells/cm^2^.

### Generation of 3D-Full Thickness Skin Model (FTSM)

FTSM were generated according to ^[42]^. Briefly, 50,000 CCD-SK1137 fibroblasts were grown in a collagen matrix for 7 days in 12-well 0.4 μm pore size ThinCerts with PET-membranes (Greiner Bio-One GmbH, Frickenhausen, Germany), which was overlayed with IMDM Medium + 10 % FBS. 5*10^5^ keratinocytes were seeded in CnT-Prime medium (CELLnTEC, Bern, Switzerland). After 3 days, medium was changed to differentiation medium (3 parts CnT-Prime 3D Barrier Medium (CELLnTEC)/ 2 parts DMEM)). On the next day, inserts were transferred to deep-well plates and lifted to the air-liquid interface (ALI) to induce keratinocyte maturation indicated by keratinization and stratification. Medium was changed every other day and models were harvested after 10 days at ALI. For barrier function studies 1 mmol Lucifer yellow (Sigma-Aldrich) was applied on top of the model for 60 min at 37°C.

### Generation of Spheroid Cocultures

SK1137 fibroblasts were seeded, with 150 µl of IMDM + 10% FBS medium, in BIOFLOAT™ 96-well plates (faCellitate, Mannheim, Germany) at a density of 5000 cells/well centrifuged at 600 rcf for 5 min (D -3). After 3 days, 100 µl of the medium were removed and keratinocytes (5000 cells/well) were added with 100 µl 3D-KGM medium (D 0) and centrifuged. 100 µl of the medium were changed on D 1 and D 3 with 3D-KGM. Coculture spheroids were collected after another 2, 4, or 6 days and processed for immunostaining.

### Immunostaining

#### FTSM

The membrane was cut out of the insert, fixed in formalin and paraffin embedded. 4 µm sections were prepared and hematoxylin and eosin (HE) stained or used for immunohistochemical staining. For immunostaining sections were de-paraffinized and antigen retrival was done in citrate buffer. Staining with the following antibodies was performed overnight at 4°C with primary antibody or concentration adjusted isotype control antibody in Signalstain buffer (CellSignaling): rabbit anti-IVL (1:1000; Abcam, ab20202), mouse anti-KRT14 (1:5000; BioRad, MCA890), rabbit anti-KRT10 (1:5000; Abcam, ab76318), rabbit anti-FLG (1:200; LSBio, LS-C751132), rabbit anti-CLD-1 (1:500; Thermofisher, 51-9000). Secondary antibodies were applied at 1:1000 for 1h (anti-mouse Cy5 labelled and anti-rabbit TxRed labelled, ThermoFisher). Sections were DAPI stained and mounted with Fluoromount G.

#### Spheroids

Spheroids were collected and fixed for 30 min at 37°C with PFA 4%. They were washed with PBS and stained with DRAQ5 (Thermofisher). After a sequential dehydration in 15 % sucrose/PBS overnight followed by 40 % sucrose/PBS for 4h at 4°C, spheroids were embedded in OTC cryomedium (Leica Biosystems) and deep-frozen. A CM-1950 cryostat (Leica Biosystems) was used to prepare 15 μm thick slices. The sections were fixed on microscope slides, washed with PBS and permeabilized with 0.1% Triton X-100 in PBS for 5 min, followed by blocking with BSA 3% in PBS for 1h at RT. Staining was performed incubating overnight at 4 °C with the following primary antibodies: rabbit anti-IVL (1:1000; Abcam, ab20202), mouse anti-KRT14 (1:1000; Thermofisher, PA5-16722), rabbit anti-KRT10 (1:1000; Abcam, ab76318), mouse anti-FLG (1:100; SantaCruz, sc-66192, AKHa1), mouse anti-CLD-1 (1:100; Thermofisher, 51-9000) diluted to the appropriate concentration in 3% BSA/PBS. After PBS washing, the slices were incubated with the following secondary antibodies (all 1:1000) for 3h at RT: donkey anti-mouse Alexa-Fluor 488 (Thermofisher, A2102), donkey anti-rabbit Alexa-Fluor 555 (Thermofisher, A32794). Nuclei were counterstained with DRAQ5 (Thermofisher, 62251). Sections were mounted with Mowiol (Roth, 0713.2).

### Histochemistry

<u>Sections</u> were stained with Mayeŕs Hämalaun/Hämatoxylin (5 min; Sigma-Aldrich, 51275) and with Eosin acid 0.5 % (5 min; Sigma-Aldrich, 1.09844) solutions, dehydrated and mounted with Entellan (Roth). Images were acquired with the upright Slide Scanner Aperio Versa 8 equipped with a Leica 20x objective 0,55/DRY and using the software Aperio Versa 1.0.4.125.

### Microscopy

FTSM - widefield: Images were acquired using a Nikon Eclipse Ci-L microscope and NIS Elements software with a 20x CFI Plan Apochromat objective at 1750×1000 pixels. Spheroids - confocal: Fluorescence images were acquired with an inverted Leica TCS SP8 confocal microscope (Leica Microsystem CSM, Mannheim, Germany) equipped with an HC APO 20x/0.75 IMM CORR objective, using the Leica Application Suite X software and Immersion Type F (Leica Microsystem, RI 1.52). All images were acquired with the same settings: resolution 1024×1024 pixels, bidirectional acquisition, scan frequency 400 Hz, line average 2x, z-step of 2 µm; laser intensity 1.2 % for 488 nm, 1 % for 555 nm and 1.7 % for 633 nm, and gain 712, 780 and 652 respectively. Live Imaging – confocal: Spheroids were loaded with CellMask Deep Red (1:1000; Thermofisher, C10046) in medium for 1 h at 37 °C, then washed 2 times isotonic buffer (329 mOsm; concentrations in mmol: KCl 5, MgCl_2_ 1, CaCl_2_ 2, HEPES 10, NaCl 145, glucose 10). To impair the cell barrier, half of the spheroids were treated with 0.1 % SDS at 37°C for 30 min. Spheroids were then placed in a µ-Slide III 3D Perfusion slide (ibidi) and fixed with Scaffolene (Gelatine nonwoven CL130 Scaffolene pads high density – crosslinked, 130 g/m^2^ from Freudenberg, perforated with a classical hole puncher). The slide was connected, using Luer-lock adaptors and tubing, to syringes containing either isotonic solution or 1 mmol LY (SIGMA, L0144). The speed of the gravity-mediated perfusion was 1.3 mL/min; complete solution exchange in the µ-well required 5-15 sec. After recording for 1.5 min with isotonic buffer perfusion, LY was applied for 3.5 min as indicated, afterward we switched again to isotonic solution perfusion. Confocal imaging used the following settings. For CellMask Deep Red: 638 nm laser, PMT 640-789, gain 650. For LY: 405 nm laser, PMT 600-600, gain 800. 3 z-planes were imaged at a depth of 0, 35, and 70 µm every 6 sec.

### Image Analysis

#### Fiji Immunostaining Analysis

For the 3D-Full Thickness model (Figure 2), the keratinocytes region was defined based on the KRT14/Lucifer Yellow merged image. Here the fibroblasts/keratinocytes boundary was defined based on KRT14 signal while the keratinocytes/air boundary was defined based on LY signal. This ROI was then applied to all the channels, the signal outside this region was set to zero and the mean intensity was measured only within the keratinocyte region. To analyze the depth-penetration of LY, a vertical ROI (12µm x 120µm) was applied to the LY image, in a representative region, and here the mean intensity was measured along the y axes. The mean intensity was plotted versus the distance from the outer border (the keratinocytes/air boundary was set as distance 0). Image intensity profiles were normalized and smoothed using a Savitzky–Golay filter to reduce noise while preserving peak shape. Peaks were automatically identified based on minimum height and prominence thresholds. For each peak, the position and height were determined, and the full width at half maximum (FWHM) was calculated to estimate peak sharpness. Linear regression was applied to the ascending and descending flanks to quantify slope characteristics. The area under the curve (AUC) was determined by trapezoidal integration across the FWHM boundaries, and a slope change point downstream of the peak was used to define the full peak width. All analyses and visualizations were implemented in Python (NumPy, SciPy, pandas, Matplotlib), and quantitative results were exported to Excel for further evaluation.

For the spheroids (Figure 4), raw confocal data were converted to multi-channel .tif files using Fiji. For each spheroid, the three-consecutive z-projections with the best signal were summed and used for the quantitative analysis. With the wand tracing tool, we manually selected the outer border of the spheroid, based on the involucrin positive signal, while the fibroblasts core region was based on the KRT14-negative region. The fibroblasts region was subtracted from the whole-spheroid region, to obtain the keratinocytes region, of which we measured the area (kerA). The keratinocyte region was applied to all the channels. The “Normalized Integrated Density” for the KRT14, KRT10, involucrin and Claudin-1 signal in the keratinocytes regions was determined as follows. By setting the lower threshold limit to 30, we defined the positive region, where we measured the area (posA) and the mean intensity (MI). We normalized the positive area to the keratinocytes area (NA=(posA/kerA) x 100) and multiplied it for the mean intensity (Normalized Integrated Density=NA x MI).

For the Lucifer Yellow Live Imaging of spheroids (Figure 8-9), the CellMask Deep Red signal was used to define the whole-spheroid region by thresholding, this ROI was then applied to the LY channel. If the Scaffolen was visible within the spheroid region, the Scaffolen signal was deleted in the LY channel, in order to avoid its contribution. At every time point, we measured the mean intensity in the spheroid region (MI_sph_) and in a smaller ROI outside the spheroids (MI_back_). MI_sph_ was normalized to the MI_back_ and was plotted over time (mean intensity as % background). For the bar histogram, the mean intensity was measured at the end of the LY perfusion (4.5 min). Every z-plane was analyzed and plotted separately (Figure 8).

For the ring analysis, only the z-plane at 35 µm was analyzed. The ImageJ erosion function was applied 30 times to the whole-spheroid ROI, to obtain an outer and an inner spheroid region. The mean intensity was measured in these two regions (MI_in_ and MI_out_), normalized (MI(%)) to the MI_back_ and plotted over time (Figure 8). In addition, the ratio between M_in_/MI_out_ was plotted over time. For the bar histogram, the MI(%) and the M_in_/MI_out_ ratio were calculated at the end of the LY perfusion (4.5 min) (Figure 9).

### Deep-Learning-based Segmentation

Nuclei images (DAPI for 3D-FT and DRAQ5 for spheroids), were pre-processed by deleting the nuclei of the fibroblast (KRT14 negative region). Nuclei in the keratinocyte’s region were then segmented by CellPose (V2.2), a deep learning-based instance segmentation tool. For each keratinocyte cell line a custom-trained model was developed as follows: at least 10 images were pre-segmented with CellPose (using the “nuclei” model) and manually corrected to generate output label masks that were then used in a “human-in-the-loop” approach to generate the custom-trained model. Polygonal ImageJ ROI files defined the overall analysis mask, as described above and in Figure 4. Cell-pose pre-computed nuclei masks were used to identify cell objects. In datasets containing a fibroblast-core mask, nuclei overlapping this region by ≥20% were discarded. To capture peri-nuclear context, a narrow zone surrounding each nucleus was generated by Euclidean distance transform and watershed segmentation, restricted to within 2 pixels of the nucleus. For each cell, mean and standard deviation of fluorescence intensity were calculated across the nuclear region and the cell zone (CZ = nucleus + peri-nuclear zone). The centroid of each nucleus was used to compute distances to structural boundaries, with distances to the spheroid hull reported as positive inside and negative outside, and analogous distances to the fibroblast-core boundary were also determined. These values were used to assign cells to three regions inside the spheroid: Region 1 (surface zone, 0–20 µm from the spheroid hull), Region 2 (inside spheroid, >20 µm from hull and not overlapping fibroblast core), Region 3 (fibroblast-adjacent, within –20 to 0 µm of the core boundary). Nuclear morphology was characterized by boundary-based estimates of major and minor axis length, aspect ratio, and eccentricity. A perpendicular axis at the centroid was compared with the shortest vector from the centroid to the spheroid surface, and the acute angle (0–90°) between these vectors was defined as the relative orientation, quantifying nuclear alignment with respect to the spheroid boundary. All analyses were performed in Python using NumPy, pandas, SciPy, scikit-image, OpenCV, tifffile, and related libraries.

### Statistics

Statistical significance was calculated with GraphPad Prism 10, as described in the figure legends. All data were tested for normal distribution by Kolmogorov-Smirnov or Shapiro Wilks normality test. The following statistical tests were applied: Two-way Anova with Multiple Comparison (corrected with Tukey-Report multiplicity adjusted P (0.05 (95% confidence interval)-Output (P value style *0.032, ** 0.0021, ***0.0002, **** <0.0001)); Ordinary One-way ANOVA; T-test unpaired (Non-Parametric Mann Whitney/Kolmogorov, Parametric-Same SD).

## AUTHOR CONTRIBUTIONS STATEMENT

*TC, CB, TF, and RR performed conceptualization. TC, NC, and VL performed investigation and methodology. TC, NC, EN, RB, and MR performed data analysis. RR and MH performed project administration and resources. TC and RR performed and wrote the original draft. RR, CB, TF, and MR edited the original draft and performed supervision. MH, RR, and TF performed funding acquisition. All authors revised the manuscript*.

## DATA AVAILABILITY STATEMENT

*The data that support the findings of this study are available from the corresponding author upon reasonable request*.

## ACKNOWLEDGMENTS

This work was funded by the German Federal Ministry of Research (BMBF) as part of the Innovation Partnership M2Aind, project DrugsData (13FH8I09IA) within the framework Starke Fachhochschulen—Impuls für die Region (FH-Impuls). This work was supported by DFG grant INST874/9-1. This work was funded by the German Federal Ministry of Education and Research (BMBF) grant 01IS21062B. We are grateful to Miriam Rittel for technical assistance.

